# Rescuing missing data in connectome-based predictive modeling

**DOI:** 10.1101/2023.06.09.544392

**Authors:** Qinghao Liang, Rongtao Jiang, Brendan D. Adkinson, Matthew Rosenblatt, Saloni Mehta, Maya L. Foster, Siyuan Dong, Chenyu You, Sahand Negahban, Harrison H. Zhou, Joseph Chang, Dustin Scheinost

**Author notes:** Qinghao Liang, Magnetic Resonance Research Center, 300 Cedar St, P.O. Box 208043, New Haven, CT 06520-8043, USA.

## Abstract

Recent evidence suggests brain-behavior predictions may require very large sample sizes. However, as the sample size increases, the amount of missing data also increases. Conventional methods, like complete-case analysis, discard useful information and shrink the sample size. To address the missing data problem, we investigated rescuing these missing data through imputation. Imputation is the substitution of estimated values for missing data to be used in downstream analyses. We integrated imputation methods into the Connectome-based Predictive Modeling (CPM) framework. Utilizing four open-source datasets—the Human Connectome Project, the Philadelphia Neurodevelopmental Cohort, the UCLA Consortium for Neuropsychiatric Phenomics, and the Healthy Brain Network (HBN)—we validated and compared our framework with different imputation methods against complete-case analysis for both missing connectomes and missing phenotypic measures scenarios. Imputing connectomes exhibited superior prediction performance on real and simulated missing data as compared to complete-case analysis. In addition, we found that imputation accuracy was a good indicator for choosing an imputation method for missing phenotypic measures but not informative for missing connectomes. In a real-world example predicting cognition using the HBN, we rescued 628 individuals through imputation, doubling the complete case sample size and increasing explained variance by 45%. Together, our results suggest that rescuing data with imputation, as opposed to discarding subjects with missing information, improves prediction performance.

## 1. Introduction

Establishing individual differences in behavior is a goal of modern neuroimaging (Dhamala et al., 2022; Dubois and Adolphs, 2016; Finn and Rosenberg, 2021; “Revisiting doubt in neuroimaging research,” 2022). However, reproducible brain-wide associations require sample sizes on the order of thousands of individuals (Genon et al., 2022; Marek et al., 2022), due to the small effect sizes of brain-behavior predictions. Using data from multiple sources increases the effect sizes of brain-behavior predictions (Dubois et al., 2018; Elliott et al., 2019; Gao et al., 2019; Jiang et al., 2020; Mellem et al., 2020; Ooi et al., 2022; Yu et al., 2020). For example, supervised learning models combining different multiple task connectomes outperform those built from a single connectome (Barron et al., 2021; Gao et al., 2019; Garrison et al., 2023), and latent variables derived from a battery of behavioral measures are more predictable as compared to a single behavioral measure (Barron et al., 2021; Dubois et al., 2018; Garrison et al., 2023; Rapuano et al., 2020). These results suggest that using connectomes and behavioral data from multiple sources is a powerful approach to characterizing brain-behavior relationships.

Nevertheless, there is a significant tradeoff when using multiple data sources of data. The amount of missing data increases. Currently, most fMRI studies only consider complete-case data, meaning that subjects with any missing behavioral or imaging data are removed from the analysis. Thus, leading to a substantial decrease in sample size. Retaining more subjects in studies is need to improve power (Genon et al., 2022; Marek et al., 2022). In addition, discarding subjects with missing data in only a single MRI modality wastes of resources, given the difficulties and cost of neuroimaging data acquisition, especially for infants, the elderly, and those with psychiatric disorders (Tejavibulya et al., 2022).

Despite being widespread in many domains of biomedicine, handling missing data is nascent in supervised learning. Most extant supervised learning studies inadequately report or handle missing data (Nijman et al., 2022). In addition, most of the literature on missing data focuses on statistical inference instead of supervised learning (Perez-Lebel et al., 2022). Only a handful of studies touch upon the theories (Josse et al., 2020) or evaluate supervised learning with missing values (Batista and Monard, 2003; Perez-Lebel et al., 2022; Poulos and Valle, 2018; Tresp et al., 1994; Zhang et al., 2010). However, the systematic evaluation of supervised learning with missing values in predictive modeling of functional connectivity data is crucial to maximizing the utility of expensive neuroimaging datasets even in the presence of missing data.

Data imputation, or the substitution of estimated values for missing data to be used in downstream analyses, is a standard way to rescue missing data. Due to the specialties of connectome data, many popular imputation methods—including statistical models like multiple imputation, maximum likelihood estimation, and fully Bayesian methods (Baraldi and Enders, 2010)—are not suitable. In connectome-based prediction analysis, the input features (edges in connectomes) are usually in a high dimension (∼30k), but the sample size is small. Even after feature selection, the dimension is still high relative to the sample size (Rosenberg et al., 2018). The connectome features are highly correlated and noisy and encode phenotypic information across many edges (Jiang et al., 2022). Moreover, the features are missed in a block-wise pattern since a whole connectome is missed. However, the redundancy in different task connectomes can be leveraged by simple imputation methods with low computational costs. Unlike connectome data imputation, most state-of-art imputation methods are suitable for the case of missing phenotypic measures.

In this work, we incorporated imputation methods into connectome-based predictive modeling (CPM) to rescue missing data in functional connectomes and behavioral measures. We investigated the performance of several imputation strategies for predicting fluid intelligence in four large open-source datasets with multiple fMRI tasks—the Human Connectome Project (HCP) (Van Essen et al., 2012), the Philadelphia Neurodevelopmental Cohort (PNC) (Poldrack et al., 2016), the UCLA Consortium for Neuropsychiatric Phenomics (CNP) (Satterthwaite et al., 2016), and the Healthy Brain Network (Alexander et al., 2017).

All four datasets contain a subset of subjects with missing fMRI data in specific tasks due to the unavailability or quality of the scans. Based on simulation analysis in HCP, we also investigated how different missingness rates of connectome data impact the performance of imputation strategies. For missing behavioral measures, we simulated missing data on ten cognitive measures and evaluated imputation methods to recover missing fluid intelligence measures. Finally, we provide a real-world example of our approach and imputed both connectomic and behavioral data in the HBN, a dataset with a a substantial missing data rate due to the difficulty in scanning children with mental health conditions. We rescued data from 628 individuals, doubling the complete case sample size and increasing explained variance by 45%. Overall, when used appropriately, data imputation is a valuable addition to neuroimaging-based prediction pipelines.

## 2. Materials and Methods

### 2.1. Datasets

Four datasets were used in our study: the Human Connectome Project (HCP) 900 Subject Release, the UCLA Consortium for Neuropsychiatric Phenomics (CNP), the Philadelphia Neurodevelopmental Cohort (PNC), and the Healthy Brain Network (HBN). The HCP dataset was used for the analysis of real missing connectomes and the simulations of missing connectomes and phenotypic measures. The CNP and PNC were used for only the analysis of real missing connectomes. The HBN dataset was used for the analysis of real missing connectomes and phenotypic measures.

#### HBN subjects

Subjects performed two naturalistic viewing sessions of movie clips from ‘Despicable Me’ and ‘The Present’ in the scanner. Ten cognitive measures were used for prediction: the Flanker Inhibitory Control and Attention (Flanker), List Sorting Working Memory (ListSort), Pattern Comparison Processing Speed (ProcSpeed), and Card Sort tests (CardSort) were selected from the NIH Toolbox; the Math Problem Solving and Numerical Operations subtests were selected from the Wechsler Individual Achievement Test (WIAT); the Fluid Reasoning, Visual Spatial Reasoning, Processing Speed, and Working Memory subtests were selected from the Wechsler Intelligence Scale for Children (WISC-V) (Alexander et al., 2017).

#### HCP subjects

Subjects performed seven distinct tasks in the scanner, including gambling, language, motor, relational, social, working memory, and emotion. The matrix reasoning test (PMAT)—a measure of fluid intelligence—was used as the phenotypic measure for prediction. For the simulations of missing phenotypic measures, we used the unadjusted score of ten cognitive measures from Dubois et al., 2018 (PicVocab, PMAT, ReadEng, VSPLOT, IWRD, PicSeq, ListSort, Flanker, CardSort, and ProcSpeed).

#### CNP subjects

Subjects performed six tasks (balloon analog risk task, paired-associative memory encoding, paired-associative memory retrieval, spatial working memory capacity, stop signal, and task switching) in the scanner. Fluid intelligence was assessed by WAIS-IV matrix reasoning (Poldrack et al., 2016).

#### PNC subjects

Subjects performed two tasks (working memory and emotion). Fluid intelligence was assessed by the 24-item version of the Penn Matrix Reasoning Test (Moore et al., 2015).

### 2.2. fMRI processing

We followed an established pipeline to create connectomic data (Barron et al., 2021). Functional images were motion corrected using SPM12. All further analyses were completed using BioImage Suites. Several covariates of no interest were regressed from the data including linear and quadratic drifts, mean cerebral-spinal-fluid (CSF) signal, mean white-matter signal, and mean gray matter signal. For additional control of possible motion-related confounds, a 24-parameter motion model (including six rigid-body motion parameters, six temporal derivatives, and these terms squared) was regressed from the data. The data were temporally smoothed with a Gaussian filter (approximate cutoff frequency = 0.12 Hz).

The brain was parcellated into 268 macroscale regions of interest using a whole-brain, functional atlas defined in a separate sample (Shen et al., 2013). For each subject, the regional time series were calculated by averaging the voxel-wise fMRI time series in a node. The pairwise Pearson correlation between all node time series was calculated and Fisher z-transformed, yielding a 268 by 268 matrix for each scan.

### 2.3. Missing data

For all datasets, connectomes were classified as missing if the imaging data was unavailable (such as subject dropouts or unshared data) or failed quality control (such as high motion or missing brain sections). Scans with a frame-to-frame displacement greater than 0.15 mm were deemed to have high motion. For analyses that included real missing data, we retained subjects who had at least one connectome, along with measures of age, sex, and fluid intelligence. For analyses with simulated missing connectomes, we used the complete-case data from the HCP dataset. Details of the HCP, CNP, and PNC datasets used in the experiments are summarized in Table 1, while the missing information for each specific task in each dataset is summarized in Fig S1. of the supplementary material.

**Table 1.** Summary of datasets (HCP, PNC, CNP) used in missing connectome data analysis.

|  |  | N | Sex (Male: Female) | Age range (Years) |
| --- | --- | --- | --- | --- |
| HCP | Complete-case | 645 | 296:349 | 22-37 |
|  | All data | 855 | 379:476 | 22-37 |
| CNP | Complete-case | 152 | 85:67 | 21-50 |
|  | All data | 244 | 139:105 | 21-50 |
| PNC | Complete-case | 682 | 310:372 | 8-21 |
|  | All data | 813 | 373:440 | 8-21 |

For analyses involving missing phenotypic data, we retained subjects who had at least one observed phenotypic variable of interest. For analyses with real missing phenotypic data, we used subjects with at least one cognitive measure from the HBN dataset. For analyses with simulated missing phenotypic data, we used subjects with complete connectome data and all ten cognitive measures from the HCP dataset. Details of the HBN and HCP datasets used in these experiments are summarized in Table 2.

**Table 2.** Summary of datasets (HBN, HCP) measures used in missing phenotypic data analysis.

|  |  | N | Sex (Male: Female) | Age range (Years) |
| --- | --- | --- | --- | --- |
| HBN | Complete-case | 698 | 447:251 | 5-21 |
|  | Complete connectome data | 893 | 575:318 | 5-22 |
|  | All data | 1326 | 851:475 | 5-22 |
| HCP | Complete-case | 514 | 240:274 | 22-37 |

### 2.4. Predictive modeling using data with missing values

To rescue data with missing connectomes, we modified Connectome-based Predictive Modeling (CPM) by incorporating imputation methods to fill in the missing values before models were trained on the completed data. Fig. 1 illustrates the missing data handling process in predictive modeling.

**Fig. 1.**
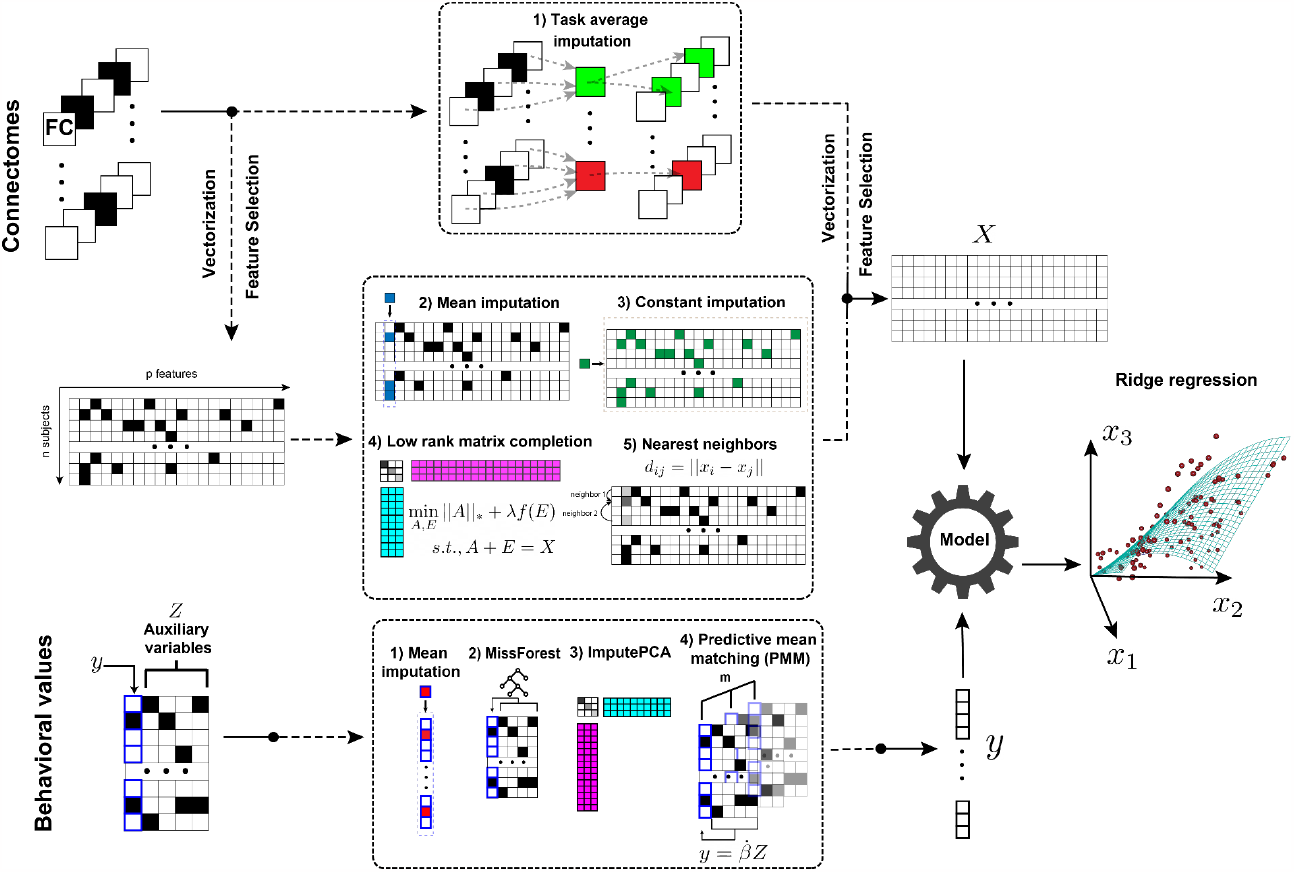
Predictive modeling framework for handling missing data. Missing connectomes are rescued using 1) Task average replacement, 2) Mean imputation, 3) Constant value imputation, 4) Robust Matrix Completion, or 5) Nearest Neighbors imputation. Missing phenotypic measures are imputed with auxiliary variables using 1) Predictive mean matching (PMM), 2) ImputePCA, 3) MissForest, or 4) Mean imputation. The phenotypic and connectivity data are then used in standardized predictive models such as ridge regression or support vector machine. The black squares represent missing data.

#### 2.4.1 Missing connectome data (missing *X*)

Five imputation methods were used to fill in missing values. Except for task average imputation, all other methods were applied on selected features to reduce the computational burden. The imputation methods are detailed below:

1. Task average imputation: For subject *i*, the mean of all observed task connectomes was calculated as *C*_*i*_. Then all missing selected features of subject *i* were replaced with the corresponding value of *C*_*i*_.
2. Mean imputation: Each missing value was replaced by the mean of the observed values along its column. Mean imputation was implemented using the python package scikit-learn (Pedregosa et al., 2011).
3. Constant values imputation: All missing values were imputed with a constant value. Here, we chose the constant value as the mean of all observed entries. Constant value imputation was implemented using the python package scikit-learn (Pedregosa et al., 2011).
4. Robust matrix completion: Given the high collinearity of edges in functional connectomes, we assume the predictive information is encoded into a lower dimensional space. Robust matrix completion recovers the low-rank matrix when part of the entries is observed and corrupted by noise (Shang et al., 2014). Robust matrix completion could be formulated as the following convex optimization problem:

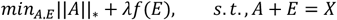

Where *X* is the corrupted input matrix (the selected features in our case), ||*A*||_*_ is the trace norm of the desired matrix *A* (output), *λ* is the parameter that controls the level of noise, *E* is the noise matrix, *f*(·) denotes the loss function. We chose the *l*_2_-norm loss in this study because the noises spread across many entries of *X*. The implementation details of this algorithm can be found in Liang et al., 2021.
5. Nearest neighbors imputation: Nearest neighbors imputation was performed by computing the Euclidean distance metric that supports missing values to find the nearest neighbors for each subject. For each missing value, values from the *k* nearest neighbors that have a value for the same feature were used to impute the missing value. The values of the neighbors were either averaged uniformly or weighted by the distance to each neighbor. If a subject had more than one missing feature, then the neighbors for that subject could be different depending on the feature being imputed. In this study, we chose *k* = 5 and used a uniform weight for each neighbor. The implementation of nearest neighbors imputation was carried out using the python package scikit-learn (Pedregosa et al., 2011).

#### 2.4.2. Missing phenotypic data (missing *y*)

When a single phenotypic measure *y* was to be predicted, we utilized a set of auxiliary variables *Z* = (*z*_1_, *z*_2_, …, *z*_*k*_) that were correlated with *y* and could also contain missing data, to impute y. When the first principal component of the variables of interest *Y* = (*y*_1_, *y*_2_, …, *y*_*k*_) was to be predicted, we imputed all missing values in *Y* and applied principal component analysis (PCA) to obtain the first principal component. The PCA coefficients were then applied to the testing set. The imputation methods are detailed below:

1. Mean imputation: Each missing value was replaced by the mean of the observed values along its column.
2. Random forest algorithm for missing data imputation (MissForest): MissForest is a non-parametric imputation method for mixed-type data. For each variable in the dataset, MissForest fits a random forest on the observed part and then predicts the missing part (the predicted values were later used in training models of other variables). The algorithm repeats these two steps until a stopping criterion (accuracy of fitting the observed part) is met. The algorithm was implemented in the R package “missForest”.
3. Regularized iterative principal component analysis (ImputePCA): Iterative Principal Component Analysis (PCA) algorithm, also known as the Expectation-maximization PCA (EMPCA) algorithm, is an expectation-maximization algorithm for a PCA fixed-effects model, where data is generated as a fixed structure having a low-rank representation corrupted by noise (Josse and Husson, 2016). Regularized iterative Principal Component Analysis uses regularized methods to tackle the overfitting problems when data is noisy, and there are many missing values. The algorithm was implemented in the function imputePCA in the R package “missMDA”.
4. Predictive mean matching (PMM): Multiple imputation by chained equations (White et al., 2011) is a robust, informative method of dealing with missing data in datasets. The procedure imputes missing data in a dataset through an iterative series of predictive models. In each iteration, each specified variable in the dataset is imputed using the other variables. These iterations should be run until it appears that convergence has been achieved. PMM selects non-missing samples with predictive values close to the predictive value of the missing sample. The closest *N* values are chosen as candidates, from which a value is chosen randomly. *M* imputation values are generated using different random initializations, and the mean value is taken to fill in the missing entries for predictive modeling. The algorithm was implemented using the R package “mice” (Buuren and Groothuis-Oudshoorn, 2011).

#### 2.4.3. Validation methods

Prediction performance of the models was evaluated using 10-fold cross-validation. Complete-case analysis served as the baseline. Subjects with missing data (*X* or *y*) were included only in the training set. The testing set consisted only of subjects with complete data. Additionally, we assessed the prediction performance of the models on subjects with missing X by randomly splitting all subjects into training and testing sets.

It is worth noting that missing data from the training and testing sets were imputed independently to prevent data leakage. A feature selection threshold of p<0.01 was used to select features in the training set. The ridge regression hyper-parameter was determined by grid searching using nested 5-fold cross-validation within the training set. For the regression task, the model performance was evaluated by the cross-validated *R*^2^, 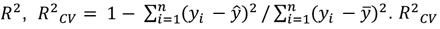 can be negative, which suggests the predictive model performs worse than simply guessing the mean of the phenotypic measure. In this case, we set *R*^3^_*CV*_ to 0 (Scheinost et al., 2019). For the classification task, the model performance was evaluated by the average of Area Under the Receiver Operating Characteristic Curve (ROC AUC) of each fold.

### 2.5. Analyses with real missing connectomes

We utilized our pipeline to analyze the HCP, CNP, and PNC datasets, and performed 100 iterations of 10-fold cross-validation to predict fluid intelligence, age, and sex. The performance of each imputation method, as well as a complete-case analysis, was calculated and reported.

### 2.6. Analyses with simulated missing connectomes

We conducted simulation tests to investigate the impact of different missingness rates and missing data mechanisms on the performance of imputation strategies. Three ypes of missingness mechanisms exist, namely missing completely at random (MCAR), missing at random (MAR), and missing not at random (MNAR) (Little and Rubin, 2019). For our experiment, we simulated two missing data scenarios. In the first scenario, we randomly deleted connectomes (MCAR). In the second scenario, we deleted connectomes with a probability proportional to the motion of the original scan, making the data missing not at random (MNAR) as the missingness was related to the data itself. We tested missingness rates ranging from 0% to 80%, averaged across the entire dataset. We performed the simulation study on the complete-case of the HCP dataset, as it has the largest amount of complete-case data. For each missingness rate, we generated 500 missingness patterns using different random seeds. For each missingness pattern, we conducted 10-fold cross-validation with a random data partition to evaluate the performance of CPM, and since the ground truth of the missing values, *x*_*true*_, was known, we also calculated the imputation accuracy of each imputation method. We assessed the accuracy of the different imputation methods using the normalized-root-mean-square error (NRMSE) between the ground truth and the imputed values *x*_*impute*_. 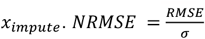, where the root-mean-square error (RMSE) is calculated as 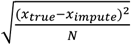 (*N* is the number of entries in *x*_*true*_) and *σ* is the standard deviation of *x*_*true*_.

### 2.7. Analyses with real missing phenotypic data

We applied our pipeline to the HBN datasets with 100 iterations of 10-fold cross-validation. Subjects with complete connectome data and missing executive function were used for predicting Flanker, ProcSpeed, and CardSort. For predicting a latent factor of cognition, we examined the performance of the models imputing cognitive measures and models imputing both cognitive measures and connectomes. Model performance of each imputation method and a complete-case study were calculated and reported.

### 2.8. Analyses with simulated missing phenotypic data

We generated 100 missingness patterns in the ten cognitive measures of the complete-case HCP data by deleting values completely at random (MCAR). Missingness rates ranged from 0 to 40%. Fluid intelligence was used as the target variable to predict, while the other nine were used as auxiliary variables. Ten iterations of CPM were run for each missingness pattern to calculate the model performance and imputation accuracy as described above.

### 2.9. Data availability statement

The data used in this study for inference and benchmarking are open-source: HCP (https://db.humanconnectome.org), CNP (https://openneuro.org/datasets/ds000030/versions/00016), and PNC (https://www.ncbi.nlm.nih.gov/projects/gap/cgi-bin/study.cgi?study_id=phs000607.v3.p2). Imputation methods are available from R or scikit-learn. The CPM code can be found at https://github.com/YaleMRRC/CPM.

## 3. Results

### 3.1. Imputing missing connectomes improves prediction performance

Using real missing data, we investigated if imputing missing connectomes improves prediction performance. Fig. 2 presents the prediction performance of phenotypic measures (sex, age, IQ) based on complete-case data and data imputed using multiple methods. Our results demonstrate that imputing missing data can significantly improve prediction accuracy compared to models built on complete data, in most cases. For sex classification, all models achieved high accuracy (>0.9) across all three datasets. Imputation improved model performance most significantly on CNP, where complete-case analysis had relatively low accuracy (0.900±0.008). However, for PNC and HCP datasets, where complete-case analysis already achieved very high accuracy, data imputation only had a marginal effect on prediction. For age and fluid intelligence (IQ) prediction, imputation led to improved performance in most cases, except for the HCP dataset. Task average imputation was found to achieve the best performance among all imputation methods. However, the relative performance of different imputation methods varies across datasets and prediction tasks.

**Fig. 2.**
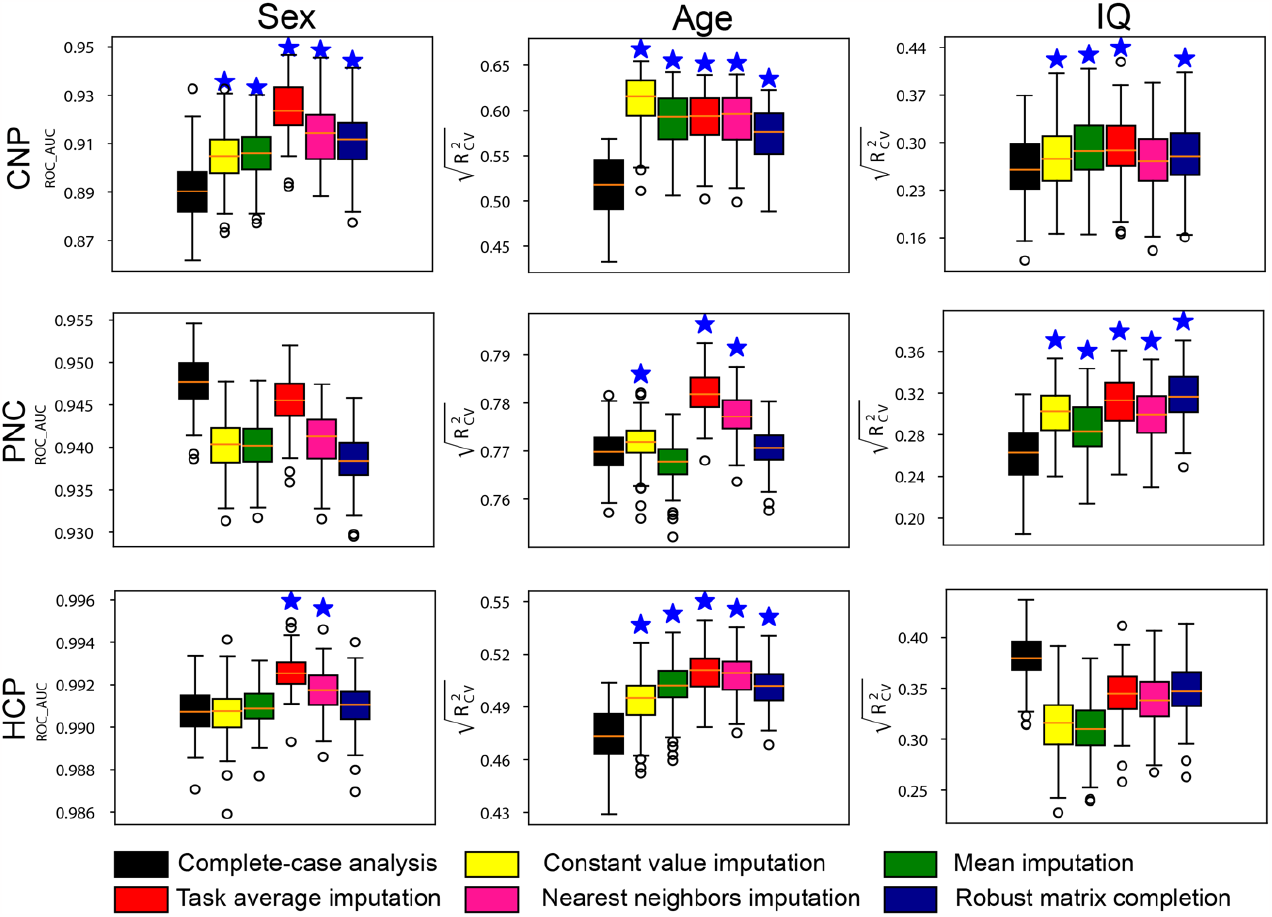
Prediction performance of models built on datasets with missing connectome data. The testing set only includes subjects with complete data. Prediction performance of sex, age and fluid intelligence based on data imputed using multiple imputation strategies in three datasets, including CNP, PNC and HCP. Stars above the boxplots indicate a significantly higher prediction performance relative to complete-case analysis (p < 0.001).

Additionally, the impact of the feature selection threshold on different models is shown in Fig. S2. Results indicated that after selecting enough features, the performance of models was relatively stable across different methods. Notably, the results presented here were derived from models where only subjects with complete data were included in the testing set. Our sensitivity analysis including subjects with incomplete data yields similar findings (Fig. S3). In some cases, the performance decline in comparison to complete-case analysis may be attributed to the disproportionate missing of critical information in the subjects added to the training set (Fig. S4).

### 3.2. Simulation to investigate the impact of missingness rates and mechanisms

We investigated the impact of missingness rates and mechanisms on imputation accuracy and prediction performance using the HCP complete-case data. As shown in Fig. 3, all imputation strategies demonstrate impaired prediction of IQ as missingness rates increase. For MCAR, task average imputation outperformed other methods at high missingness rates, while all methods had a similar performance at low rates. For MNAR, robust matrix completion had the highest performance, while task average imputation performed the worst. There was no significant difference in terms of imputation accuracy between the two missing mechanisms. The accuracy of imputation methods in recovering connectome data decreased with increasing missingness rates. Notably, task average imputation showed the worst performance in recovering connectome data. Robust matrix completion was the best method for recovering connectome data for missingness rates below 0.4. However, it showed impaired performance for missingness rates above 0.4.

**Fig. 3.**
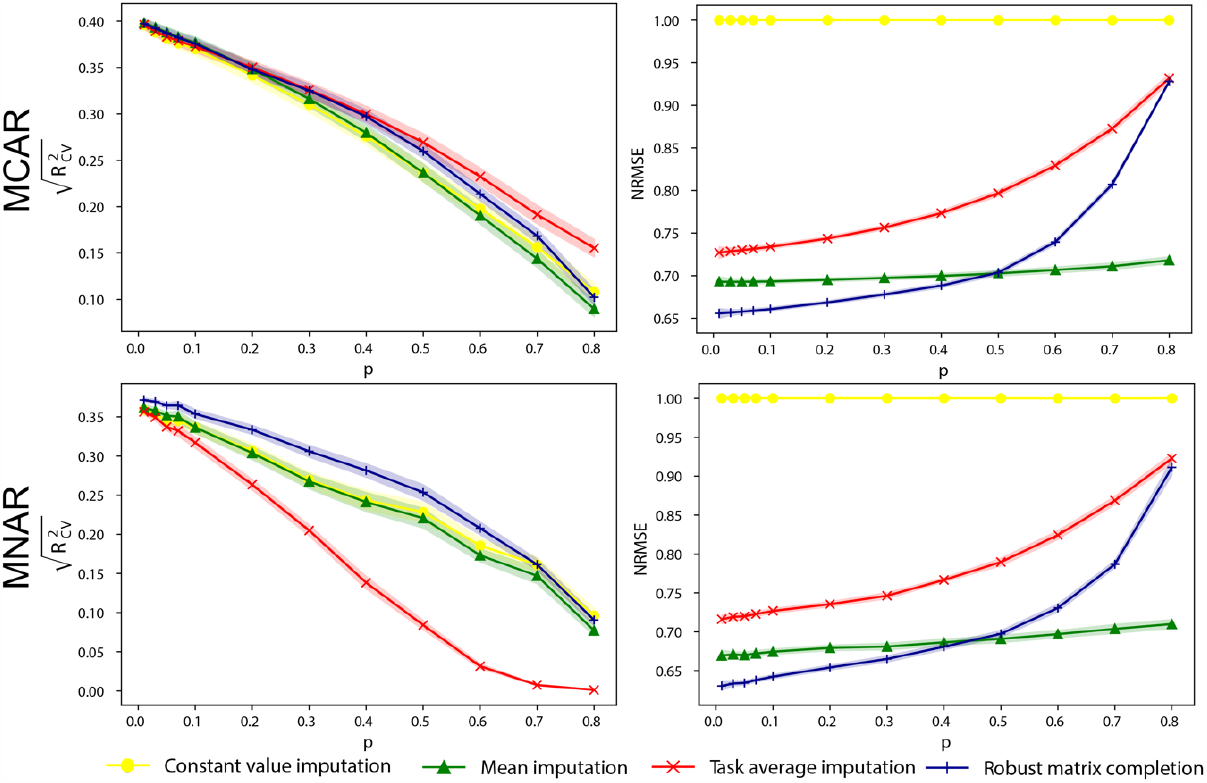
Prediction performance of models built with subjects of simulated missing connectomes. Prediction performance 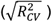 of models predicting IQ using different missing data handling strategies as a function of missingness rates *p*. Imputation accuracy (NRMSE) as a function of missingness rates *p*. The shadow areas were 95% confidence intervals. The missing data were generated on the complete data in HCP through random sampling (MCAR) or sampling.

### 3.3. Imputing missing phenotypic data improves prediction performance

In the HBN dataset, a substantial number of subjects have missing connectomes and cognitive measures. Through imputation, we were able to rescue a number of subjects: 33 for CardSort, 34 for Flanker, and 42 for ProcSpeed. For predicting the first principal component of ten cognitive measures, 195 subjects were rescued by imputing the missing cognitive measures. Only subjects with missing data were included in the training set, and the prediction performance was evaluated on the same testing set used for complete-case analysis.

As depicted in Fig. 4a and b, imputing missing cognitive measures significantly improved the prediction performance compared to complete-case analysis. Among the four imputation methods used, mean imputation yielded the lowest prediction performance.

**Fig. 4.**
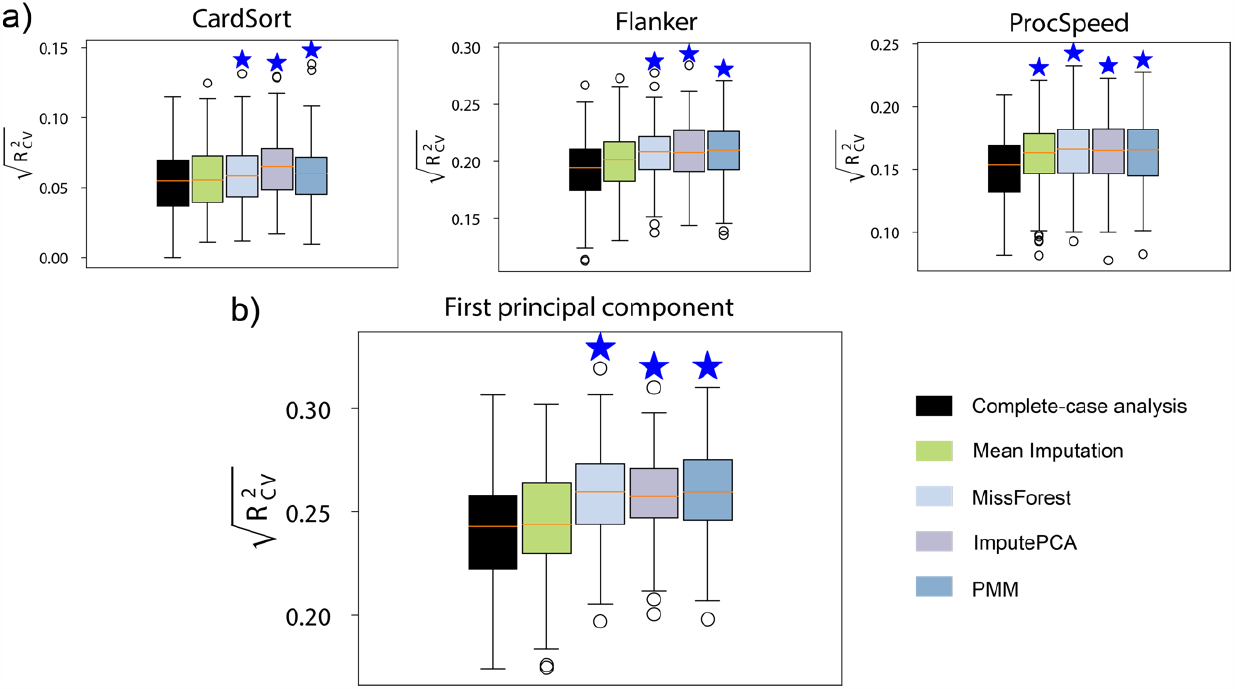
Prediction performance of models built with HBN subjects with missing phenotypic measures. a) Performance of predicting a single cognitive measure. b) Performance of predicting the first principal component of ten cognitive measures.

### 3.4. Simulation to investigate the impact of missingness rates in phenotypic measures

Using the IQ data from the HCP, we conducted a simulation analysis to investigate the impact of missingness rates on prediction performance. We observed a decrease in prediction performance for all models as the missingness rate increased (Fig. 5). However, three imputation methods, namely MissForest, ImputePCA, and PMM, outperformed the complete-case analysis data, while mean imputation harmed prediction performance. Notably, ImputePCA consistently achieved the highest imputation accuracy across all missingness rates, followed by PMM and MissForest, whereas mean imputation had the lowest imputation accuracy (Fig. 5). Unlike missing connectomes, we observed that methods with higher imputation accuracy achieved better prediction performance simultaneously for missing cognitive measures.

**Fig. 5.**
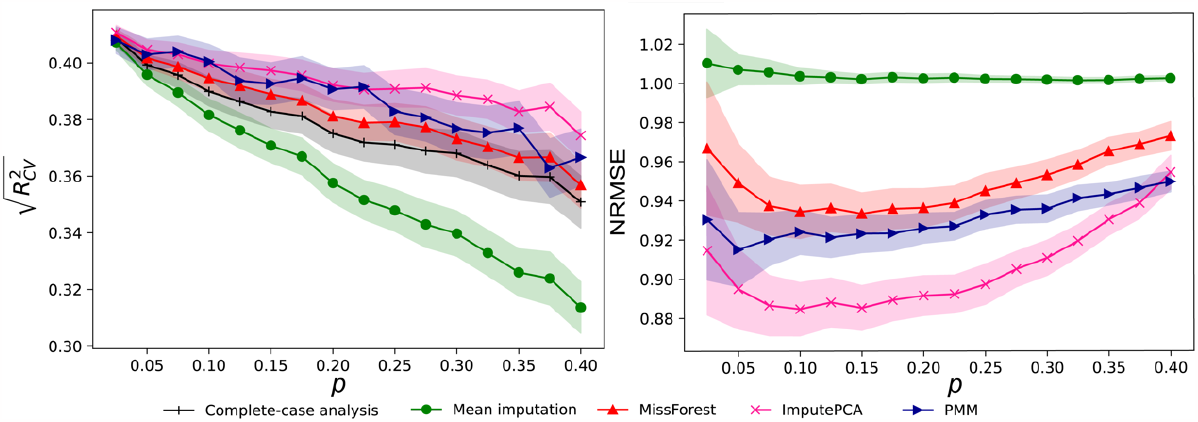
Prediction performance of models built with subjects of simulated missing phenotypic measures. Prediction performance 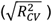 of models using different missing data handling strategies as a function of missingness rates *p*. Imputation accuracy (NRMSE) as a function of missingness rates *p*. The shadow areas were 95% confidence intervals.

### 3.5. Imputing missing connectomes and missing phenotypic data to rescue the maximum amount of data

Finally, we present a real-world example of the benefits of data imputation in predicting cognition in the HBN dataset. In this example, we predicted the first principal component of ten cognitive measures. The prediction performance in the complete case was 0.295±0.024. 628 subjects were rescued by imputing both the missing cognitive measures and connectomes. For imputing missing connectome data, we employed two simple methods: mean imputation and task average imputation. However, due to its inferior performance, meaning imputation was not used for imputing missing cognitive measures. The prediction performance was significantly greater when including the imputed data in the training set (Fig. 6). The combination of task average imputation and MissForest achieved the highest prediction performance (0.347±0.018). Overall, correlation between observed and predicted was ∼20% greater and explained variance was ∼45% greater after including imputed data in the training set.

**Fig. 6.**
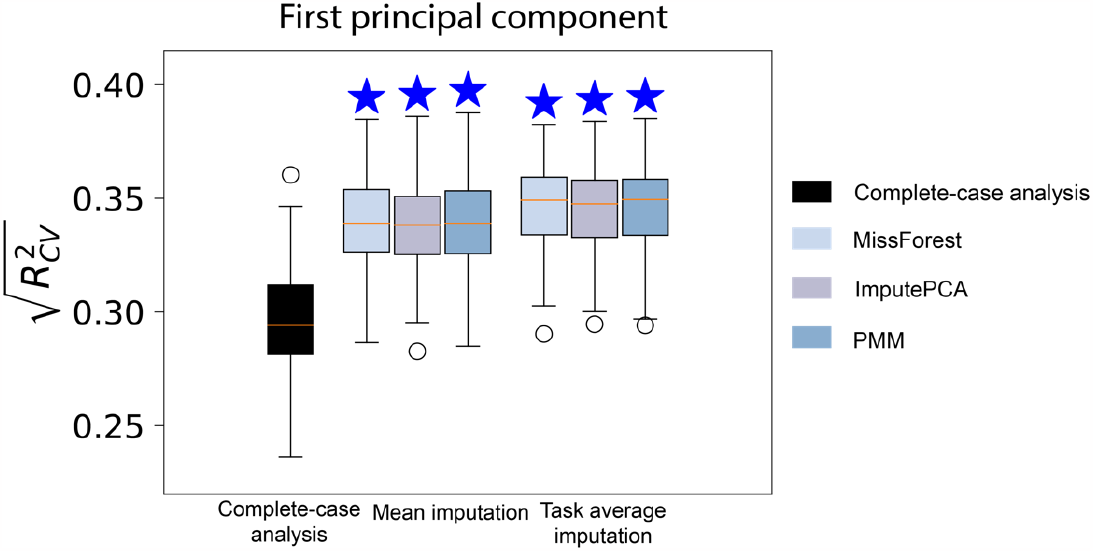
Prediction performance of models built with HBN subjects with both missing connectomes and missing phenotypic measures. Missing connectomes were imputed using mean imputation and task average imputation. Missing cognitive measures were imputed using MissForest, ImputePCA, and PMM. In this example, we predicted the first principal component of ten cognitive measures from the HBN dataset.

## 4. Discussion

In this study, we investigated rescuing missing connectomes and behavioral measures in connectome-based predictive modeling using data imputation. Rescued, imputed data were used to increase the size of the training data. For missing connectomes, we evaluated the effects of data imputation for predicting sex, age, fluid intelligence, and executive function on three independent data sets with missing connectomes. Results show that imputing missing data improved prediction performance over complete-case analysis. Moreover, simulation analyses showed that imputation strategies that best recover missing data do not necessarily signify better prediction performance, and the missing data mechanism affects the performance of different imputation methods differently. For missing phenotypic measures, improved performance over complete-case analysis was also achieved by most imputation methods. Contrary to missing connectomes, imputation strategies with higher accuracy achieved better model performance. In a real-world example, we show that including imputed connectomes and cognitive data to increase training sample sizes can significantly increase prediction performance for difficult to collect samples. Overall, our results show the potential of imputation methods to address incomplete data for predictive modeling.

### 4.1. Data imputation improves prediction performance

The contribution of our study is demonstrating that rescuing missing data with imputation significantly improves model performance. In the case of missing connectomes, most models built with incomplete data outperformed complete-case analysis. The improvement in prediction performance scales with the proportion of data rescued. In the CNP and HBN datasets, the prediction performance showed a more significant increase compared to PNC and HCP, as the rescued data is relatively larger compared to the complete data. However, the gains of including more subjects will be saturated when the prediction performance reaches a certain level. Notably, simple imputation methods such as task average imputation achieved the highest prediction performance in real missing data analysis. However, when connectomes are deleted with a probability proportional to motion, task average imputation failed to work. This result is because subjects with high motion tend to lose more connectomes, which hinders the efficacy of imputation with the average across tasks. Our results have implications for connectome-based prediction research that integrates multi-source information from distinct fMRI tasks. We showed that even leveraging simple imputation strategies can significantly improve prediction performance. A possible explanation for this result is that the intrinsic network structure, which is task-independent, contributes the most predictive information (Cole et al., 2014; Fox et al., 2005; Geerligs et al., 2015; Mill et al., 2017; Vincent et al., 2007). Moreover, recent years have witnessed the increasing availability of large-scale open data cohorts such as UK Biobank (Sudlow et al., 2015) and the Adolescent Brain Cognitive Development (ABCD) (Karcher and Barch, 2021), which collected multimodal MRI data from tens of thousands of participants. In this context, applying imputation strategies can facilitate the use of neuroimaging data to the greatest extent possible and further accelerate the pace of establishing robust and generalizable prediction models.

Robust matrix completion, which works by representing the original high-dimensional data using low-rank approximation, ranks highest among all tested methods in PNC and second in HCP and CNP for predicting fluid intelligence. This result lends support to the existence of a low dimensional space across cognitive tasks (intrinsic brain information) that is representative of the general information encoded in distinct fMRI tasks but highly informative in characterizing individual differences in cognition and behavior (Breakspear, 2017; Fox and Raichle, 2007; Gao et al., 2021; Krienen et al., 2014; Mennes et al., 2013; Shine et al., 2019).

All methods except mean imputation, which diminishes individual differences, outperformed complete-case analysis in imputing missing behavioral measures. The auxiliary variables used in the experiments are all cognitive ability measures that are correlated with each other (Dubois et al., 2018). The effectiveness of using multiple cognitive measures accords with the hypothesis that a general factor of intelligence could be derived from several cognitive tasks through factor analysis (de la Fuente et al., 2021). It is also worth noting some similar observations in meta-matching methods (He et al., 2022), which translate predictive models from large-scale datasets to new unseen phenotypes in small-scale studies. In meta-matching, prediction improvements were driven by correlations between training and test meta-set phenotypes. In our case, the imputation accuracy of the target variable—in other words, how well the target variable could be predicted by auxiliary variables—was highly correlated with prediction improvements. The similarity is not surprising given that both methods aim to include subjects with phenotypes correlated with the target phenotype for model training.

### 4.2. Higher imputation accuracy does not guarantee higher prediction performance

Imputation accuracy, the accuracy of estimating missing data, was generally considered the gold standard for evaluating imputation methods (Lin and Tsai, 2020). However, our results indicate that imputation strategies that can effectively estimate missing data do not necessarily signify better prediction performance. Interestingly, our missing connectome simulation achieved the highest prediction accuracy using the average connectome across tasks. Nevertheless, this imputation strategy had the worst performance in recovering connectome data. These findings gain support from other studies investigating the relationship between imputation accuracy and prediction performance (Perez-Lebel et al., 2022). They can be partially attributed to the fact that the information recovered by imputation methods is not the most informative for prediction. In the case of connectomes with low signal-to-noise ratios (Vizioli et al., 2021), the methods with high imputation accuracy might recover noise that harms the modeling building. However, in the case of missing behavioral values (missing *y*), higher imputation accuracy corresponds to higher prediction performance because noisy labels severely degrade the generalization performance of machine learning models, especially when the sample size is small (Song et al., 2022).

### 4.3. Limitations and future directions

Several potential limitations should be acknowledged when interpreting the current findings. First, we could not draw any clear conclusion about which imputation strategy is best for recovering incomplete data for prediction. Indeed, given the complexities of the problem, each method may only work for some conditions. However, our study provides preliminary insight that even a simple imputation strategy can significantly improve prediction performance and open up opportunities for future studies to integrate other imputation methods into connectome-based prediction. Second, the impact of missingness mechanisms was not considered in this pipeline. In neuroimaging studies, data are usually not missing completely at random. The imputation process could also lead to bias toward a certain subgroup of subjects (Groenwold and Dekkers, 2020; Jakobsen et al., 2017). For example, head motion can sometimes predict certain phenotypes (Zeng et al., 2014), in which case the missingness indicator provides predictive modeling information (Donders et al., 2006). A crucial next step for research will be examining the potential impact of missing data mechanisms and imputation methods on datasets with larger sample sizes like the UK Biobank and ABCD. Third, the current study was performed in a functional connectome with a high-dimension and multicollinear nature. It remains unclear to what extent the imputation strategies work for other imaging modalities like structural MRI and DTI. In the next step of our research, imputation methods for other imaging modalities will be developed and examined. Finally, in neuroimaging studies, missing phenotypic measures are less common since the cost of labeling data is much lower than that of acquiring imaging data. The missingness could be caused by combining data from different studies. Typically, cases with imputed *y* contain no information about the regression of *y* on *X* (Von Hippel, 2007). Imputation is only worthwhile when there are auxiliary variables available.

## 5. Conclusion

In summary, we proposed a framework for CPM that rescues missing values in the imaging data and behavioral measures with imputation techniques. The proposed method enables us to harness the information of samples with missing data and achieve higher prediction performance compared to complete-case analysis. Moreover, we aim to raise awareness of the missing data problem in the neuroimaging community. Overall, our results suggest that subjects with incomplete data are valuable for predictive modeling in neuroimaging studies which are usually limited by the sample size.

## Author contributions

Dustin Scheinost and Qinghao Liang conceptualized the study; Qinghao Liang performed the data analysis; Dustin Scheinost, Rongtao Jiang, and Qinghao Liang wrote the paper. All authors contributed to the interpretation and discussion of the results.

## Acknowledgements

Data were provided in part by the Human Connectome Project, WU-Minn Consortium (Principal Investigators: David Van Essen and Kamil Ugurbil; 1U54MH091657) funded by the 16 NIH Institutes and Centers that support the NIH Blueprint for Neuroscience Research; and by the McDonnell Center for Systems Neuroscience at Washington University. The second part of the data used in this study was supported by the Consortium for Neuropsychiatric Phenomics (NIH Roadmap for Medical Research grants UL1-DE019580, RL1MH083268, RL1MH083269, RL1DA024853, RL1MH083270, RL1LM009833, PL1MH083271, and PL1NS062410). Support for the collection of the data for the Philadelphia Neurodevelopmental Cohort (PNC) was provided by grant RC2MH089983, awarded to Raquel Gur, and RC2MH089924, awarded to Hakon Hakonarson. Subjects were recruited and genotyped through the Center for Applied Genomics (CAG) at The Children’s Hospital in Philadelphia (CHOP). Phenotypic data collection occurred at the CAG/CHOP and at the Brain Behavior Laboratory, University of Pennsylvania.

## Supplementary materials

**Fig. S1.**
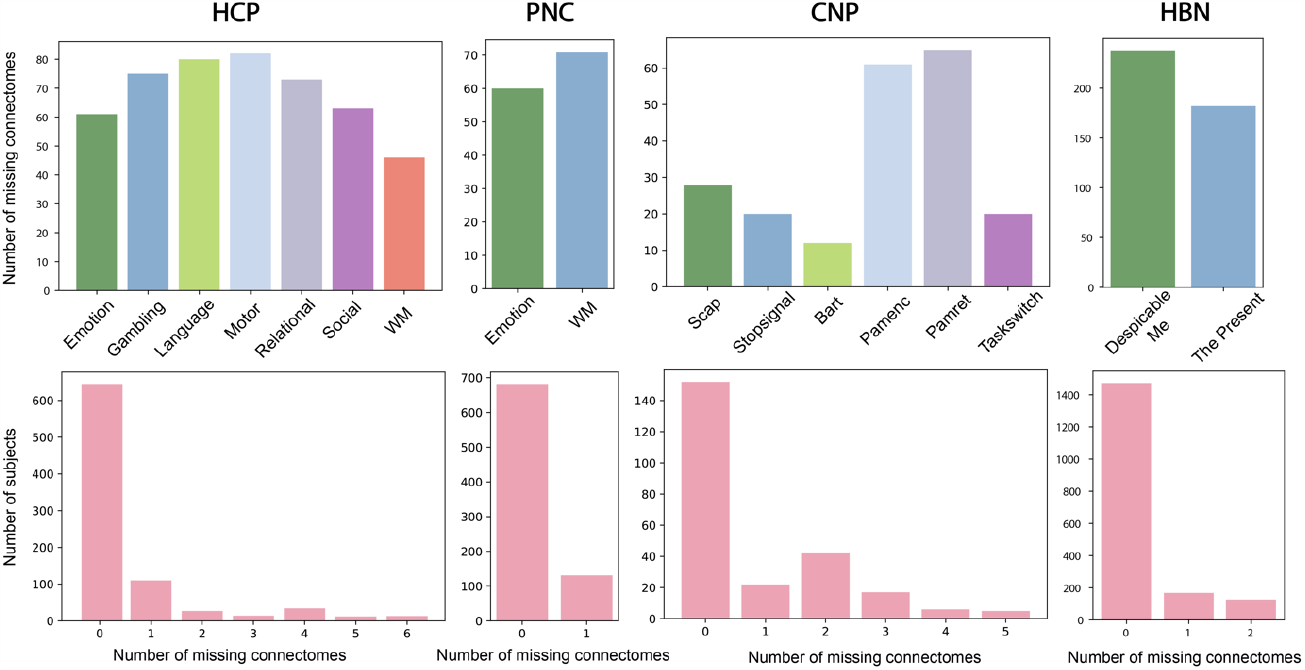
Missingness of connectomes in the experimental datasets (HCP, PNC, CNP, HBN). The first row shows the number of subjects that had missing connectome data for each task. The second row summarizes the counts of subjects by the number of missing connectomes.

**Fig. S2.**
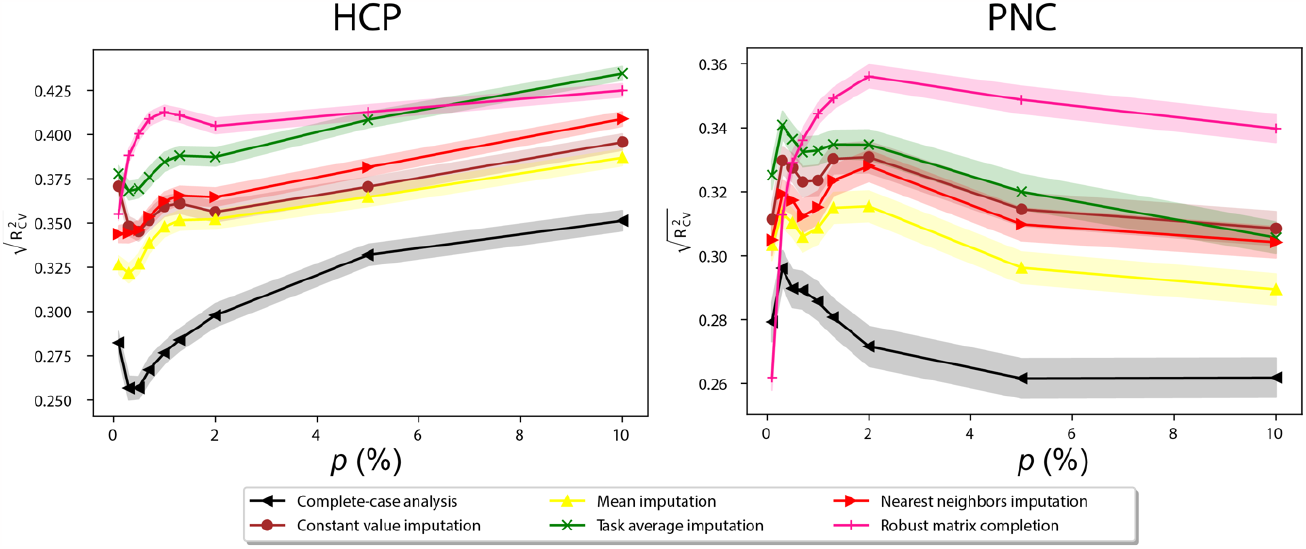
Prediction performance of IQ models built with subjects of missing connectomes given different feature selection percentages.

**Fig. S3.**
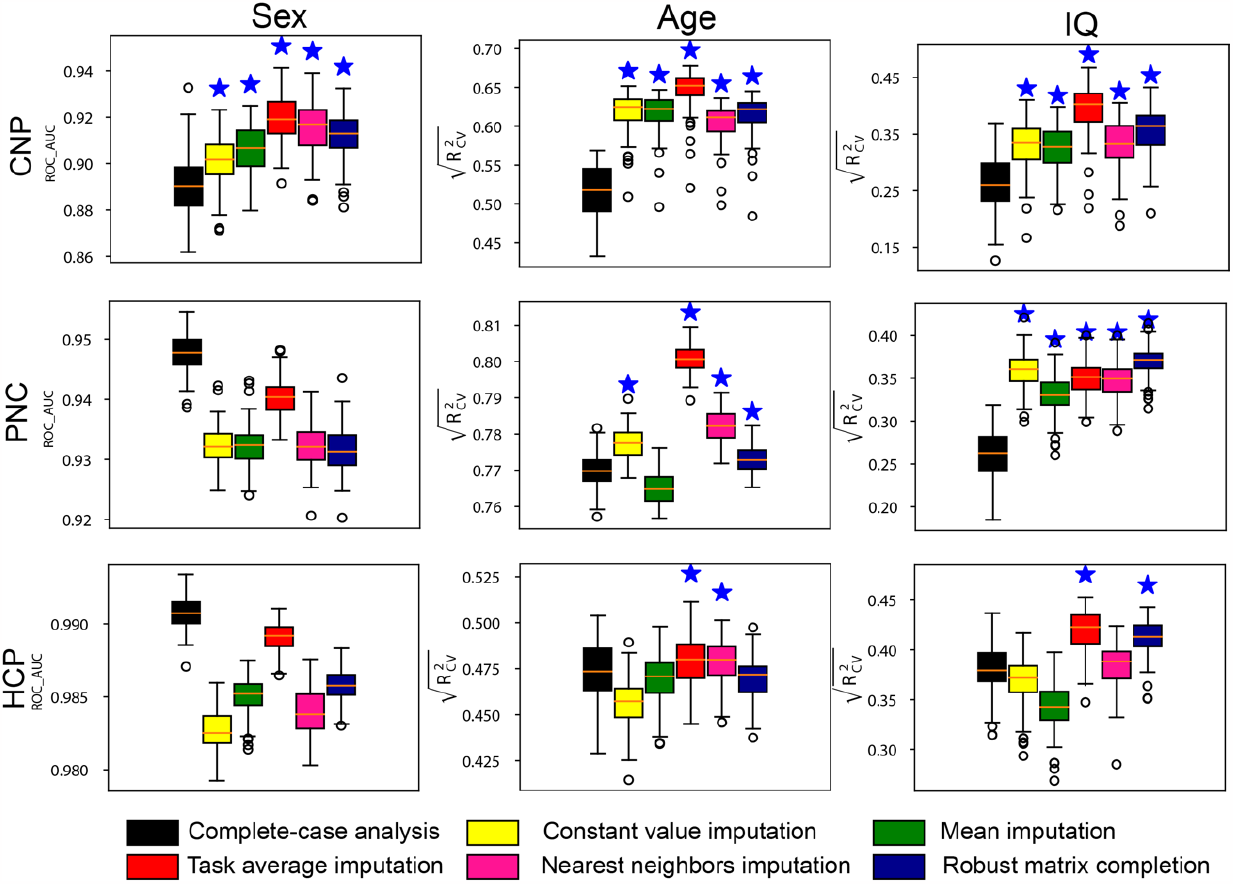
Prediction performance of models built on datasets with missing connectome data. The testing set includes subjects with missing connectome data. Prediction performance of sex, age and fluid intelligence based on data imputed using multiple imputation strategies in three datasets, including CNP, PNC and HCP. Stars above the boxplots indicate a significantly higher prediction performance relative to complete-case analysis (p < 0.001).

**Fig. S4.**
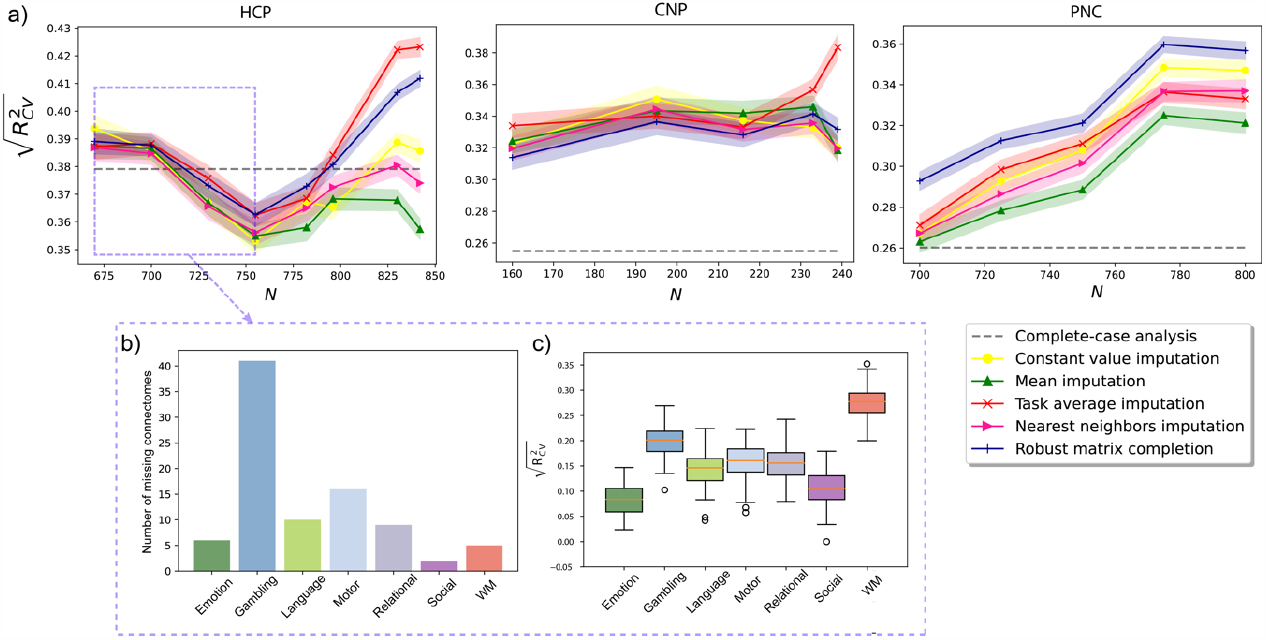
Effect of including more subjects with higher missingness on prediction performance (validated using all data). We iteratively added subjects with higher missing connectome rates to investigate the impact of including subjects with different missingness rates. a) The plots show the prediction performance as the function of the number of subjects. Subjects with a higher rate of missing connectomes were incrementally added to the complete-case data. The shaded areas were 95% confidence intervals. To investigate the reason for the initial prediction performance decline in HCP dataset, we showed: b) The missingness pattern for subjects who missed one connectome. The bars show the counts of missing connectomes of each task. c) Prediction performance for fluid intelligence using connectome data from each single fMRI task in HCP shown in boxplots. The model was built using the complete-case data.

## Reference

Alexander, L.M., Escalera, J., Ai, L., Andreotti, C., Febre, K., Mangone, A., Vega-Potler, N., Langer, N., Alexander, A., Kovacs, M., Litke, S., O’Hagan, B., Andersen, J., Bronstein, B., Bui, A., Bushey, M., Butler, H., Castagna, V., Camacho, N., Chan, E., Citera, D., Clucas, J., Cohen, S., Dufek, S., Eaves, M., Fradera, B., Gardner, J., Grant-Villegas, N., Green, G., Gregory, C., Hart, E., Harris, S., Horton, M., Kahn, D., Kabotyanski, K., Karmel, B., Kelly, S.P., Kleinman, K., Koo, B., Kramer, E., Lennon, E., Lord, C., Mantello, G., Margolis, A., Merikangas, K.R., Milham, J., Minniti, G., Neuhaus, R., Levine, A., Osman, Y., Parra, L.C., Pugh, K.R., Racanello, A., Restrepo, A., Saltzman, T., Septimus, B., Tobe, R., Waltz, R., Williams, A., Yeo, A., Castellanos, F.X., Klein, A., Paus, T., Leventhal, B.L., Craddock, R.C., Koplewicz, H.S., Milham, M.P., 2017. An open resource for transdiagnostic research in pediatric mental health and learning disorders. Sci. Data 4, 170181. https://doi.org/10.1038/sdata.2017.181

Baraldi, A.N., Enders, C.K., 2010. An introduction to modern missing data analyses. J. Sch. Psychol. 48, 5–37. https://doi.org/10.1016/j.jsp.2009.10.001

Barron, D.S., Gao, S., Dadashkarimi, J., Greene, A.S., Spann, M.N., Noble, S., Lake, E.M.R., Krystal, J.H., Constable, R.T., Scheinost, D., 2021. Transdiagnostic, Connectome-Based Prediction of Memory Constructs Across Psychiatric Disorders. Cereb. Cortex 31, 2523–2533. https://doi.org/10.1093/cercor/bhaa371

Batista, G.E.A.P.A., Monard, M.C., 2003. An analysis of four missing data treatment methods for supervised learning. Appl. Artif. Intell. 17, 519–533. https://doi.org/10.1080/713827181

Breakspear, M., 2017. Dynamic models of large-scale brain activity. Nat. Neurosci. 20, 340–352. https://doi.org/10.1038/nn.4497

Buuren, S. van, Groothuis-Oudshoorn, K., 2011. mice: Multivariate Imputation by Chained Equations in R. J. Stat. Softw. 45, 1–67. https://doi.org/10.18637/jss.v045.i03

Cole, M.W., Bassett, D.S., Power, J.D., Braver, T.S., Petersen, S.E., 2014. Intrinsic and Task-Evoked Network Architectures of the Human Brain. Neuron 83, 238–251. https://doi.org/10.1016/j.neuron.2014.05.014

de la Fuente, J., Davies, G., Grotzinger, A.D., Tucker-Drob, E.M., Deary, I.J., 2021. A general dimension of genetic sharing across diverse cognitive traits inferred from molecular data. Nat. Hum. Behav. 5, 49–58. https://doi.org/10.1038/s41562-020-00936-2

Dhamala, E., Yeo, B.T.T., Holmes, A.J., 2022. One Size Does Not Fit All: Methodological Considerations for Brain-Based Predictive Modeling in Psychiatry. Biol. Psychiatry. https://doi.org/10.1016/j.biopsych.2022.09.024

Donders, A.R.T., van der Heijden, G.J.M.G., Stijnen, T., Moons, K.G.M., 2006. Review: A gentle introduction to imputation of missing values. J. Clin. Epidemiol. 59, 1087–1091. https://doi.org/10.1016/j.jclinepi.2006.01.014

Dubois, J., Adolphs, R., 2016. Building a Science of Individual Differences from fMRI. Trends Cogn. Sci. 20, 425–443. https://doi.org/10.1016/j.tics.2016.03.014

Dubois, J., Galdi, P., Paul, L.K., Adolphs, R., 2018. A distributed brain network predicts general intelligence from resting-state human neuroimaging data. Philos. Trans. R. Soc. B Biol. Sci. 373, 20170284. https://doi.org/10.1098/rstb.2017.0284

Elliott, M.L., Knodt, A.R., Cooke, M., Kim, M.J., Melzer, T.R., Keenan, R., Ireland, D., Ramrakha, S., Poulton, R., Caspi, A., Moffitt, T.E., Hariri, A.R., 2019. General functional connectivity: Shared features of resting-state and task fMRI drive reliable and heritable individual differences in functional brain networks. NeuroImage 189, 516–532. https://doi.org/10.1016/j.neuroimage.2019.01.068

Finn, E.S., Rosenberg, M.D., 2021. Beyond fingerprinting: Choosing predictive connectomes over reliable connectomes. NeuroImage 239, 118254. https://doi.org/10.1016/j.neuroimage.2021.118254

Fox, M.D., Raichle, M.E., 2007. Spontaneous fluctuations in brain activity observed with functional magnetic resonance imaging. Nat. Rev. Neurosci. 8, 700–711. https://doi.org/10.1038/nrn2201

Fox, M.D., Snyder, A.Z., Vincent, J.L., Corbetta, M., Van Essen, D.C., Raichle, M.E., 2005. The human brain is intrinsically organized into dynamic, anticorrelated functional networks. Proc. Natl. Acad. Sci. 102, 9673–9678. https://doi.org/10.1073/pnas.0504136102

Gao, S., Greene, A.S., Constable, R.T., Scheinost, D., 2019. Combining multiple connectomes improves predictive modeling of phenotypic measures. NeuroImage 201, 116038. https://doi.org/10.1016/j.neuroimage.2019.116038

Gao, S., Mishne, G., Scheinost, D., 2021. Nonlinear manifold learning in functional magnetic resonance imaging uncovers a low-dimensional space of brain dynamics. Hum. Brain Mapp. 42, 4510–4524. https://doi.org/10.1002/hbm.25561

Garrison, K.A., Sinha, R., Potenza, M.N., Gao, S., Liang, Q., Lacadie, C., Scheinost, D., 2023. Transdiagnostic Connectome-Based Prediction of Craving. Am. J. Psychiatry 180, 445–453. https://doi.org/10.1176/appi.ajp.21121207

Geerligs, L., Rubinov, M., Cam-CAN Henson, R.N., 2015. State and Trait Components of Functional Connectivity: Individual Differences Vary with Mental State. J. Neurosci. 35, 13949–13961. https://doi.org/10.1523/JNEUROSCI.1324-15.2015

Genon, S., Eickhoff, S.B., Kharabian, S., 2022. Linking interindividual variability in brain structure to behaviour. Nat. Rev. Neurosci. 23, 307–318. https://doi.org/10.1038/s41583-022-00584-7

Groenwold, R.H.H., Dekkers, O.M., 2020. Missing data: the impact of what is not there. Eur. J. Endocrinol. 183, E7–E9. https://doi.org/10.1530/EJE-20-0732

He, T., An, L., Chen, P., Chen, J., Feng, J., Bzdok, D., Holmes, A.J., Eickhoff, S.B., Yeo, B.T.T., 2022. Meta-matching as a simple framework to translate phenotypic predictive models from big to small data. Nat. Neurosci. 25, 795–804. https://doi.org/10.1038/s41593-022-01059-9

Jakobsen, J.C., Gluud, C., Wetterslev, J., Winkel, P., 2017. When and how should multiple imputation be used for handling missing data in randomised clinical trials – a practical guide with flowcharts. BMC Med. Res. Methodol. 17, 162. https://doi.org/10.1186/s12874-017-0442-1

Jiang, R., Woo, C.-W., Qi, S., Wu, J., Sui, J., 2022. Interpreting Brain Biomarkers: Challenges and solutions in interpreting machine learning-based predictive neuroimaging. IEEE Signal Process. Mag. 39, 107–118. https://doi.org/10.1109/MSP.2022.3155951

Jiang, R., Zuo, N., Ford, J.M., Qi, S., Zhi, D., Zhuo, C., Xu, Y., Fu, Z., Bustillo, J., Turner, J.A., Calhoun, V.D., Sui, J., 2020. Task-induced brain connectivity promotes the detection of individual differences in brain-behavior relationships. NeuroImage 207, 116370. https://doi.org/10.1016/j.neuroimage.2019.116370

Josse, J., Husson, F., 2016. missMDA: A Package for Handling Missing Values in Multivariate Data Analysis. J. Stat. Softw. 70, 1–31. https://doi.org/10.18637/jss.v070.i01

Josse, J., Prost, N., Scornet, E., Varoquaux, G., 2020. On the consistency of supervised learning with missing values. https://doi.org/10.48550/arXiv.1902.06931

Karcher, N.R., Barch, D.M., 2021. The ABCD study: understanding the development of risk for mental and physical health outcomes. Neuropsychopharmacology 46, 131–142. https://doi.org/10.1038/s41386-020-0736-6

Krienen, F.M., Yeo, B.T.T., Buckner, R.L., 2014. Reconfigurable task-dependent functional coupling modes cluster around a core functional architecture. Philos. Trans. R. Soc. B Biol. Sci. 369, 20130526. https://doi.org/10.1098/rstb.2013.0526

Liang, Q., Negahban, S., Chang, J., Zhou, H.H., Scheinost, D., 2021. Connectome-Based Predictive Modelling With Missing Connectivity Data Using Robust Matrix Completion, in: 2021 IEEE 18th International Symposium on Biomedical Imaging (ISBI). Presented at the 2021 IEEE 18th International Symposium on Biomedical Imaging (ISBI), pp. 738–742. https://doi.org/10.1109/ISBI48211.2021.9434138

Lin, W.-C., Tsai, C.-F., 2020. Missing value imputation: a review and analysis of the literature (2006–2017). Artif. Intell. Rev. 53, 1487–1509. https://doi.org/10.1007/s10462-019-09709-4

Little, R.J.A., Rubin, D.B.R., 2019. Statistical Analysis with Missing Data, Third Edition.

Marek, S., Tervo-Clemmens, B., Calabro, F.J., Montez, D.F., Kay, B.P., Hatoum, A.S., Donohue, M.R., Foran, W., Miller, R.L., Hendrickson, T.J., Malone, S.M., Kandala, S., Feczko, E., Miranda-Dominguez, O., Graham, A.M., Earl, E.A., Perrone, A.J., Cordova, M., Doyle, O., Moore, L.A., Conan, G.M., Uriarte, J., Snider, K., Lynch, B.J., Wilgenbusch, J.C., Pengo, T., Tam, A., Chen, J., Newbold, D.J., Zheng, A., Seider, N.A., Van, A.N., Metoki, A., Chauvin, R.J., Laumann, T.O., Greene, D.J., Petersen, S.E., Garavan, H., Thompson, W.K., Nichols, T.E., Yeo, B.T.T., Barch, D.M., Luna, B., Fair, D.A., Dosenbach, N.U.F., 2022. Reproducible brain-wide association studies require thousands of individuals. Nature 603, 654–660. https://doi.org/10.1038/s41586-022-04492-9

Mellem, M.S., Liu, Y., Gonzalez, H., Kollada, M., Martin, W.J., Ahammad, P., 2020. Machine Learning Models Identify Multimodal Measurements Highly Predictive of Transdiagnostic Symptom Severity for Mood, Anhedonia, and Anxiety. Biol. Psychiatry Cogn. Neurosci. Neuroimaging 5, 56–67. https://doi.org/10.1016/j.bpsc.2019.07.007

Mennes, M., Kelly, C., Colcombe, S., Castellanos, F.X., Milham, M.P., 2013. The Extrinsic and Intrinsic Functional Architectures of the Human Brain Are Not Equivalent. Cereb. Cortex 23, 223–229. https://doi.org/10.1093/cercor/bhs010

Mill, R.D., Ito, T., Cole, M.W., 2017. From connectome to cognition: The search for mechanism in human functional brain networks. NeuroImage, Functional Architecture of the Brain 160, 124–139. https://doi.org/10.1016/j.neuroimage.2017.01.060

Moore, T.M., Reise, S.P., Gur, R.E., Hakonarson, H., Gur, R.C., 2015. Psychometric properties of the Penn Computerized Neurocognitive Battery. Neuropsychology 29, 235–246. https://doi.org/10.1037/neu0000093

Nijman, S., Leeuwenberg, A., Beekers, I., Verkouter, I., Jacobs, J., Bots, M., Asselbergs, F., Moons, K., Debray, T., 2022. Missing data is poorly handled and reported in prediction model studies using machine learning: a literature review. J. Clin. Epidemiol. 142, 218–229. https://doi.org/10.1016/j.jclinepi.2021.11.023

Ooi, L.Q.R., Chen, J., Zhang, S., Kong, R., Tam, A., Li, J., Dhamala, E., Zhou, J.H., Holmes, A.J., Yeo, B.T.T., 2022. Comparison of individualized behavioral predictions across anatomical, diffusion and functional connectivity MRI. NeuroImage 263, 119636. https://doi.org/10.1016/j.neuroimage.2022.119636

Pedregosa, F., Varoquaux, G., Gramfort, A., Michel, V., Thirion, B., Grisel, O., Blondel, M., Prettenhofer, P., Weiss, R., Dubourg, V., Vanderplas, J., Passos, A., Cournapeau, D., Brucher, M., Perrot, M., Duchesnay, É., 2011. Scikit-learn: Machine Learning in Python. J. Mach. Learn. Res. 12, 2825–2830.

Perez-Lebel, A., Varoquaux, G., Le Morvan, M., Josse, J., Poline, J.-B., 2022. Benchmarking missing-values approaches for predictive models on health databases. GigaScience 11, giac013. https://doi.org/10.1093/gigascience/giac013

Poldrack, R.A., Congdon, E., Triplett, W., Gorgolewski, K.J., Karlsgodt, K.H., Mumford, J.A., Sabb, F.W., Freimer, N.B., London, E.D., Cannon, T.D., Bilder, R.M., 2016. A phenome-wide examination of neural and cognitive function. Sci. Data 3, 160110. https://doi.org/10.1038/sdata.2016.110

Poulos, J., Valle, R., 2018. Missing Data Imputation for Supervised Learning. Appl. Artif. Intell. 32, 186–196. https://doi.org/10.1080/08839514.2018.1448143

Rapuano, K.M., Rosenberg, M.D., Maza, M.T., Dennis, N.J., Dorji, M., Greene, A.S., Horien, C., Scheinost, D., Todd Constable, R., Casey, B.J., 2020. Behavioral and brain signatures of substance use vulnerability in childhood. Dev. Cogn. Neurosci. 46, 100878. https://doi.org/10.1016/j.dcn.2020.100878

Revisiting doubt in neuroimaging research, 2022.. Nat. Neurosci. 25, 833–834. https://doi.org/10.1038/s41593-022-01125-2

Rosenberg, M.D., Casey, B.J., Holmes, A.J., 2018. Prediction complements explanation in understanding the developing brain. Nat. Commun. 9, 589. https://doi.org/10.1038/s41467-018-02887-9

Satterthwaite, T.D., Connolly, J.J., Ruparel, K., Calkins, M.E., Jackson, C., Elliott, M.A., Roalf, D.R., Hopson, R., Prabhakaran, K., Behr, M., Qiu, H., Mentch, F.D., Chiavacci, R., Sleiman, P.M.A., Gur, R.C., Hakonarson, H., Gur, R.E., 2016. The Philadelphia Neurodevelopmental Cohort: A publicly available resource for the study of normal and abnormal brain development in youth. NeuroImage, Sharing the wealth: Brain Imaging Repositories in 2015 124, 1115–1119. https://doi.org/10.1016/j.neuroimage.2015.03.056

Scheinost, D., Noble, S., Horien, C., Greene, A.S., Lake, E.MR., Salehi, M., Gao, S., Shen, X., O’Connor, D., Barron, D.S., Yip, S.W., Rosenberg, M.D., Constable, R.T., 2019. Ten simple rules for predictive modeling of individual differences in neuroimaging. NeuroImage 193, 35–45. https://doi.org/10.1016/j.neuroimage.2019.02.057

Shang, F., Liu, Y., Cheng, J., Cheng, H., 2014. Robust Principal Component Analysis with Missing Data, in: Proceedings of the 23rd ACM International Conference on Conference on Information and Knowledge Management, CIKM ‘14. Association for Computing Machinery, New York, NY, USA, pp. 1149–1158. https://doi.org/10.1145/2661829.2662083

Shen, X., Tokoglu, F., Papademetris, X., Constable, R.T., 2013. Groupwise whole-brain parcellation from resting-state fMRI data for network node identification. NeuroImage 82, 403–415. https://doi.org/10.1016/j.neuroimage.2013.05.081

Shine, J.M., Breakspear, M., Bell, P.T., Ehgoetz Martens, K.A., Shine, R., Koyejo, O., Sporns, O., Poldrack, R.A., 2019. Human cognition involves the dynamic integration of neural activity and neuromodulatory systems. Nat. Neurosci. 22, 289–296. https://doi.org/10.1038/s41593-018-0312-0

Song, H., Kim, M., Park, D., Shin, Y., Lee, J.-G., 2022. Learning From Noisy Labels With Deep Neural Networks: A Survey. IEEE Trans. Neural Netw. Learn. Syst. 1–19. https://doi.org/10.1109/TNNLS.2022.3152527

Sudlow, C., Gallacher, J., Allen, N., Beral, V., Burton, P., Danesh, J., Downey, P., Elliott, P., Green, J., Landray, M., Liu, B., Matthews, P., Ong, G., Pell, J., Silman, A., Young, A., Sprosen, T., Peakman, T., Collins, R., 2015. UK Biobank: An Open Access Resource for Identifying the Causes of a Wide Range of Complex Diseases of Middle and Old Age. PLOS Med. 12, e1001779. https://doi.org/10.1371/journal.pmed.1001779

Tejavibulya, L., Rolison, M., Gao, S., Liang, Q., Peterson, H., Dadashkarimi, J., Farruggia, M.C., Hahn, C.A., Noble, S., Lichenstein, S.D., Pollatou, A., Dufford, A.J., Scheinost, D., 2022. Predicting the future of neuroimaging predictive models in mental health. Mol. Psychiatry 27, 3129–3137. https://doi.org/10.1038/s41380-022-01635-2

Tresp, V., Neuneier, R., Ahmad, S., 1994. Efficient Methods for Dealing with Missing Data in Supervised Learning, in: Advances in Neural Information Processing Systems. MIT Press.

Van Essen, D.C., Ugurbil, K., Auerbach, E., Barch, D., Behrens, T.E.J., Bucholz, R., Chang, A., Chen, L., Corbetta, M., Curtiss, S.W., Della Penna, S., Feinberg, D., Glasser, M.F., Harel, N., Heath, A.C., Larson-Prior, L., Marcus, D., Michalareas, G., Moeller, S., Oostenveld, R., Petersen, S.E., Prior, F., Schlaggar, B.L., Smith, S.M., Snyder, A.Z., Xu, J., Yacoub, E., WU-Minn HCP Consortium, 2012. The Human Connectome Project: a data acquisition perspective. NeuroImage 62, 2222–2231. https://doi.org/10.1016/j.neuroimage.2012.02.018

Vincent, J.L., Patel, G.H., Fox, M.D., Snyder, A.Z., Baker, J.T., Van Essen, D.C., Zempel, J.M., Snyder, L.H., Corbetta, M., Raichle, M.E., 2007. Intrinsic functional architecture in the anaesthetized monkey brain. Nature 447, 83–86. https://doi.org/10.1038/nature05758

Vizioli, L., Moeller, S., Dowdle, L., Akçakaya, M., De Martino, F., Yacoub, E., Uğurbil, K., 2021. Lowering the thermal noise barrier in functional brain mapping with magnetic resonance imaging. Nat. Commun. 12, 5181. https://doi.org/10.1038/s41467-021-25431-8

Von Hippel, P.T., 2007. Regression with Missing Ys: An Improved Strategy for Analyzing Multiply Imputed Data. Sociol. Methodol. 37, 83–117. https://doi.org/10.1111/j.1467-9531.2007.00180.x

White, I.R., Royston, P., Wood, A.M., 2011. Multiple imputation using chained equations: Issues and guidance for practice. Stat. Med. 30, 377–399. https://doi.org/10.1002/sim.4067

Yu, J., Rawtaer, I., Fam, J., Feng, L., Kua, E.-H., Mahendran, R., 2020. The individualized prediction of cognitive test scores in mild cognitive impairment using structural and functional connectivity features. NeuroImage 223, 117310. https://doi.org/10.1016/j.neuroimage.2020.117310

Zeng, L.-L., Wang, D., Fox, M.D., Sabuncu, M., Hu, D., Ge, M., Buckner, R.L., Liu, H., 2014. Neurobiological basis of head motion in brain imaging. Proc. Natl. Acad. Sci. 111, 6058–6062. https://doi.org/10.1073/pnas.1317424111

Zhang, S., Wu, X., Zhu, M., 2010. Efficient missing data imputation for supervised learning, in: 9th IEEE International Conference on Cognitive Informatics (ICCI’10). Presented at the 9th IEEE International Conference on Cognitive Informatics (ICCI’10), pp. 672–679. https://doi.org/10.1109/COGINF.2010.5599826

